# An accessible, efficient and global approach for the large-scale sequencing of bacterial genomes

**DOI:** 10.1101/2020.07.22.200840

**Authors:** Blanca M. Perez-Sepulveda, Darren Heavens, Caisey V. Pulford, Alexander V. Predeus, Ross Low, Hermione Webster, Christian Schudoma, Will Rowe, James Lipscombe, Chris Watkins, Benjamin Kumwenda, Neil Shearer, Karl Costigan, Kate S. Baker, Nicholas A. Feasey, Jay C. D. Hinton, Neil Hall, The 10KSG consortium

## Abstract

We have developed an efficient and inexpensive pipeline for streamlining large-scale collection and genome sequencing of bacterial isolates. Evaluation of this method involved a worldwide research collaboration focused on the model organism *Salmonella enterica*, the 10KSG consortium. By optimising a logistics pipeline that collected isolates as thermolysates, permitting shipment in ambient conditions, the project assembled a diverse collection of 10,419 clinical and environmental isolates from low- and middle-income countries in less than one year. The bacteria were obtained from 51 countries/territories dating from 1949 to 2017, with a focus on Africa and Latin-America. All isolates were collected in barcoded tubes and genome sequenced using an optimised DNA extraction method and the LITE pipeline for library construction. After Illumina sequencing, the total reagent cost was less than USD$10 per genome. Our method can be applied to genome-sequence other large bacterial collections at a relatively low cost, within a limited timeframe, to support global collaborations.

## Introduction

Whole genome sequencing (WGS) is an important tool that has revolutionised our understanding of bacterial disease over the past decade^1–4^. Recognising the immense advantages that WGS data provides for surveillance, functional genomics and population dynamics, both public health and research communities have adopted genome-based approaches.

Until recently, large-scale bacterial genome projects could only be performed in a handful of sequencing centres around the world. Here, we aimed to make this technology accessible to bacterial laboratories worldwide. The high demand for sequencing human genomes has driven down the costs of sequencing reagents to below USD$1,000 per sample^5–7^. However, the genome sequencing of thousands of microorganisms has remained expensive due to costs associated with sample transportation and library construction.

The number of projects focused on sequencing the genomes of collections of key pathogens has increased markedly over recent years. Whilst the first *Vibrio cholerae* next-generation WGS study was based on 23 genomes^8^, a recent study involved 1,070 isolates from 45 African countries^9^ and identified the origin of the most recent cholera pandemic. *Mycobacterium tuberculosis,* another major human pathogen, was originally sequenced on the 100-isolate scale in 2010^10^, whilst recent publications used 3,651^11^ or 10,209^12^ genomes to evaluate the accuracy of antibiotic resistance prediction. Other successful large-scale next-generation WGS projects for pathogens include *Salmonella, Shigella, Staphylococcus,* and pneumococcus *(Streptococcus pneumoniae*)^13–16^.

One of the most significant challenges facing scientific researchers in low- and middle-income (LMI) countries is the streamlining of surveillance with scientific collaborations. For a combination of reasons, the regions associated with the greatest burden of severe bacterial disease have inadequate access to WGS technology and usually have to rely on expensive and bureaucratic processes for sample transport and sequencing. This has prevented the adoption of large-scale genome sequencing and analysis of bacterial pathogens for public health and surveillance in LMI countries^17^. Here, we have established an efficient and relatively inexpensive pipeline for the worldwide collection and sequencing of bacterial genomes. To evaluate our pipeline, we used the model organism *Salmonella enterica,* a pathogen with a global significance^18^.

Non-typhoidal *Salmonella* (NTS) are widely associated with enterocolitis in humans, a zoonotic disease that is linked to the industrialisation of food production. Because of the scale of human cases of enterocolitis and concerns related to food safety, more genome sequences have been generated for *Salmonella* than for any other genus. The number of publicaly available sequenced *Salmonella* genomes will soon reach 300,000^19^, and are available from several public repositories such as the European Nucleotide Archive (ENA, https://www.ebi.ac.uk/ena), the Sequence Read Archive (SRA, https://www.ncbi.nlm.nih.gov/sra), and Enterobase (https://enterobase.warwick.ac.uk/species/index/senterica). However, there has been limited genome-based surveillance of foodborne infections in LMI countries, and the available genomic dataset does not accurately represent the *Salmonella* pathogens that are currently causing disease across the world.

In recent years, new lineages of NTS serovars Typhimurium and Enteritidis have been recognised as common causes of invasive bloodstream infections (iNTS disease), responsible for about 77,000 deaths per year worldwide^20^. Approximately 80% of deaths due to iNTS disease occurs in sub-Saharan Africa, where iNTS disease has become endemic^21^. The new *Salmonella* lineages responsible for bloodstream infections of immunocompromised individuals are characterised by genomic degradation, altered prophage repertoires and novel multidrug resistant plasmids^22, 23^.

We saw a need to simplify and expand genome-based surveillance of salmonellae from Africa and other parts of the world, involving isolates associated with invasive disease and gastroenteritis in humans, and extended to bacteria derived from animals and the environment. We optimised a pipeline for streamlining the large-scale collection and sequencing of samples from LMI countries with the aim of facilitating access to WGS and worldwide collaboration. Our pipeline represents a relatively inexpensive and robust tool for the generation of bacterial genomic data from LMI countries, allowing investigation of the epidemiology, drug resistance and virulence factors of isolates.

## Results

### Development of an optimised logistics pipeline

The “10,000 *Salmonella* genomes” (10KSG; https://10k-salmonella-genomes.com/) is a global consortium that includes collaborators from 25 institutions and a variety of settings, including research and reference laboratories across 16 countries. Limited funding resources prompted us to design an approach that ensured accurate sample tracking and captured comprehensive metadata for individual bacterial isolates whilst minimising costs for the consortium. A key driver was to assemble a set of genomic data that would be as informative and robust as possible.

Members of the 10KSG provided access to 10,419 bacterial isolates from collections that spanned 51 LMI countries and regions (such as Reunion Island, an overseas department and region of the French Republic). We optimised the logistics of specimen collection and the transport of materials to the sequencing centre in the UK. The standardised protocols for metadata and sample submission were coordinated in three different languages (English, French and Spanish), which facilitated collaboration across several countries (Fig. 1).

**Fig. 1.**
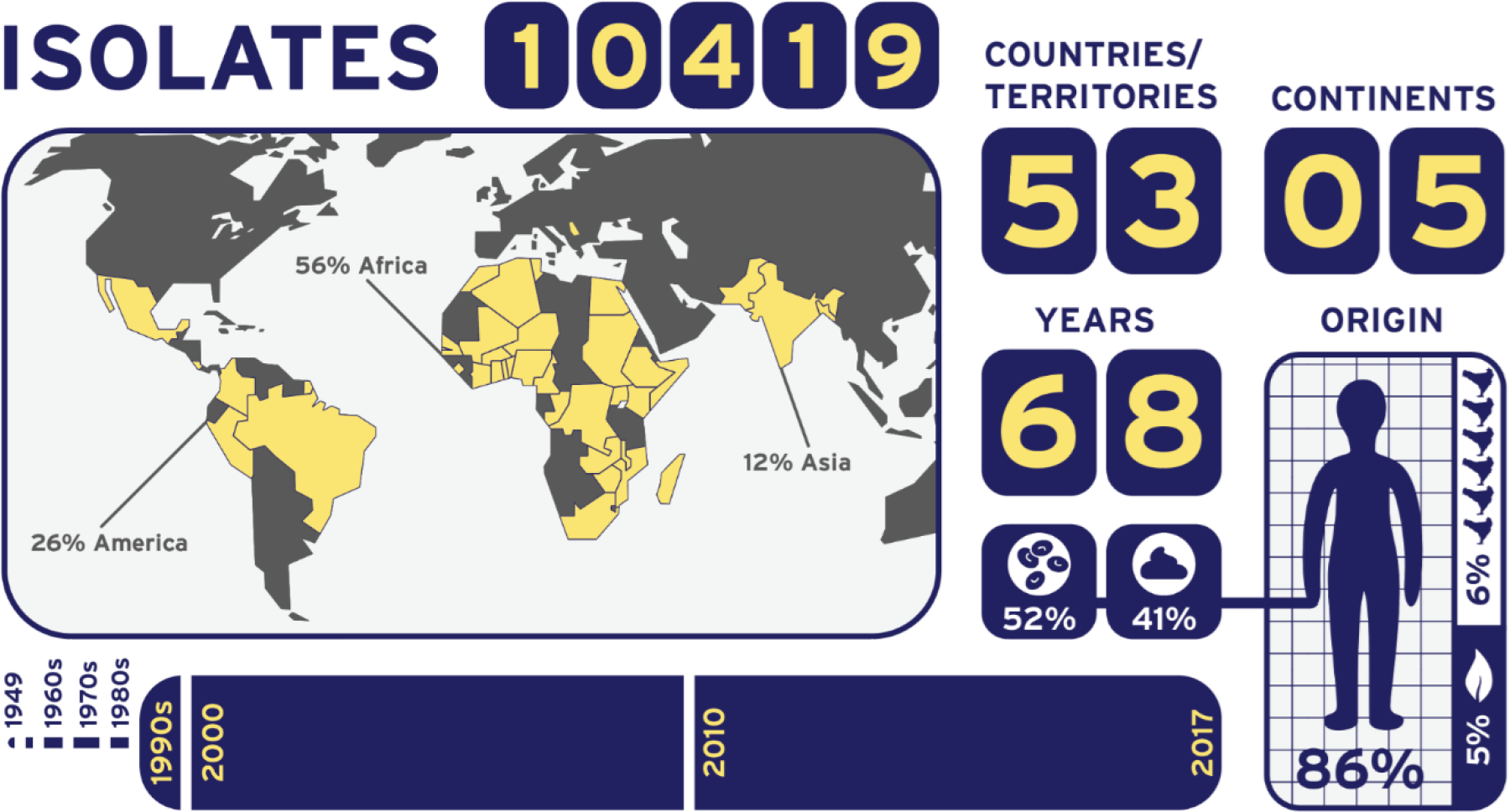
Summary of the geographical origin, timeline and body site source of 10,419 bacterial isolates. The 10,419 isolates were collected from 53 countries/territories spanning 5 continents (America, Africa, Asia, Europe, & Oceania), with most isolates originating from Africa (56%) and America (26%). The samples were mostly of human origin (86%), of which 52% were blood isolates, 41% were stool isolates, and 7% from other body compartments. About 5% samples originated from environmental sources, 6% were of animal origin, and 3% unknown. The bacterial pathogens were isolated over a 68-year time period, from 1949 to 2017. The majority of samples were isolated after 1990.

A crucial criterion for inclusion of *Salmonella* isolates in this study was the availability of detailed metadata and phenotypic information, to maximise the insights that could be generated from bacterial genomics. We created a standardised metadata table for input of relevant parameters. The metadata template was divided into categories, including unique sample identifier, date of isolation, geographical location, source niche (human, animal or environmental isolate) & type (body compartment). We also collected data regarding the antimicrobial susceptibility of isolates, and captured additional information related to individual studies. We created a unified metadata master-form (Supplementary Table 1) by manual concatenation and curation of individual metadata forms.

### Development of thermolysates and sample collection

The main challenges for the global collection of bacterial samples are temperature-control and biological safety during transport. As refrigerated logistic chains are expensive, shipments should be at ambient temperature to minimise costs. To ensure biosafety, it was important to avoid the accidental transport of hazard group three (HG3) isolates (e.g., *S*. Typhi and *S*. Paratyphi A)^24^. Accordingly, we optimised a protocol for production of “thermolysates” that inactivated bacterial cells and permitted ambient temperature transport and adherence to containment level two (CL2) laboratory regulations, coupled with effective genomic DNA extraction for WGS (Supplementary Table 2). Inactivation of *Salmonella* can be achieved at temperatures between 55°C to 70°C for as little as 15 s at high temperature (≥ 95°C)^25^. We optimised the method for generation of “thermolysates” by inactivating bacterial cultures at high temperature (95°C for 20 min). The optimisation involved testing under three different temperatures (90°C, 95°C or 100°C) and different incubation times (10 and 20 min). We also tested the effective inactivation of other non-*Salmonella* Gram-positive (*Staphylococcus aureus*) and Gram-negative (*Escherichia coli*) organisms (Supplementary Table 2).

Temperature is a key factor in the transportation of samples, especially in some LMI countries where dry ice is expensive and difficult to source, and access to international courier companies is limited or very costly. To allow transport without refrigeration, we tested the stability of the resulting thermolysates at room temperature for more than seven days by controlling the quality of extracted DNA (Supplementary Table 2). Minimising the steps required for sample collection allowed us to reach collaborators with limited access to facilities and personnel.

We collected samples using screwed-cap barcoded tubes (FluidX tri-coded jacket 0.7 mL, Brooks Life Sciences, 68-0702-11) costing USD$0.23 each, which we distributed from the UK to collaborators worldwide. Individually barcoded tubes were organised in FluidX plates in a 96-well format, each with their own barcode. Both QR codes and human-readable barcodes were included on each tube to ensure that the correct samples were always sequenced, and to permit the replacement of individual tubes when required.

All isolates were obtained in compliance with the Nagoya protocol^26^. The combination of method optimisation, development and distribution of easy-to-follow protocols in English and Spanish (French was used only for communication), generating thermolysates and using barcoded tubes, the process of collecting the bacterial isolates was completed within one year. Barcoded tubes were distributed to collaborators, including an extra ~20% to permit replacements as required. In total, 11,823 tubes were used in the study, of which 10,419 were returned to the sequencing centre containing bacterial thermolysates for DNA extraction and genome sequencing. A comprehensive list of isolates is available in Supplementary Table 3. To validate this approach for bacteria other than *Salmonella,* ~25% (2,573, 24.7%) of the samples were isolates from a variety of genera, including Gram-negatives such as *Shigella* and *Klebsiella,* and Gram-positives such as *Staphylococcus.*

### DNA extraction, library construction, quality control and genome sequencing

Our high-throughput DNA extraction and library construction pipeline was designed to be versatile, scalable and robust, capable of processing thousands of samples in a time and cost-efficient manner. The procedure included DNA extraction, quality control (QC), normalisation, sequencing library construction, pooling, size selection and sequencing. The time taken for each step, and the associated consumable cost, is shown in Table 1. All the parts of the pipeline are scalable and can be run simultaneously with robots, allowing hundreds of samples to be processed each day, in a 96-well format. With dedicated pre- and post-PCR robots, up to 768 bacterial samples were processed each day. The total consumable cost for extraction of DNA and genome sequence generation was less than USD$10 per sample (excluding staff time). Given the high-throughput nature of this project, and the difficulty in optimising the processes to account for every possible variation in DNA/library quality and quantity, this cost includes a 20% contingency.

**Table 1.**
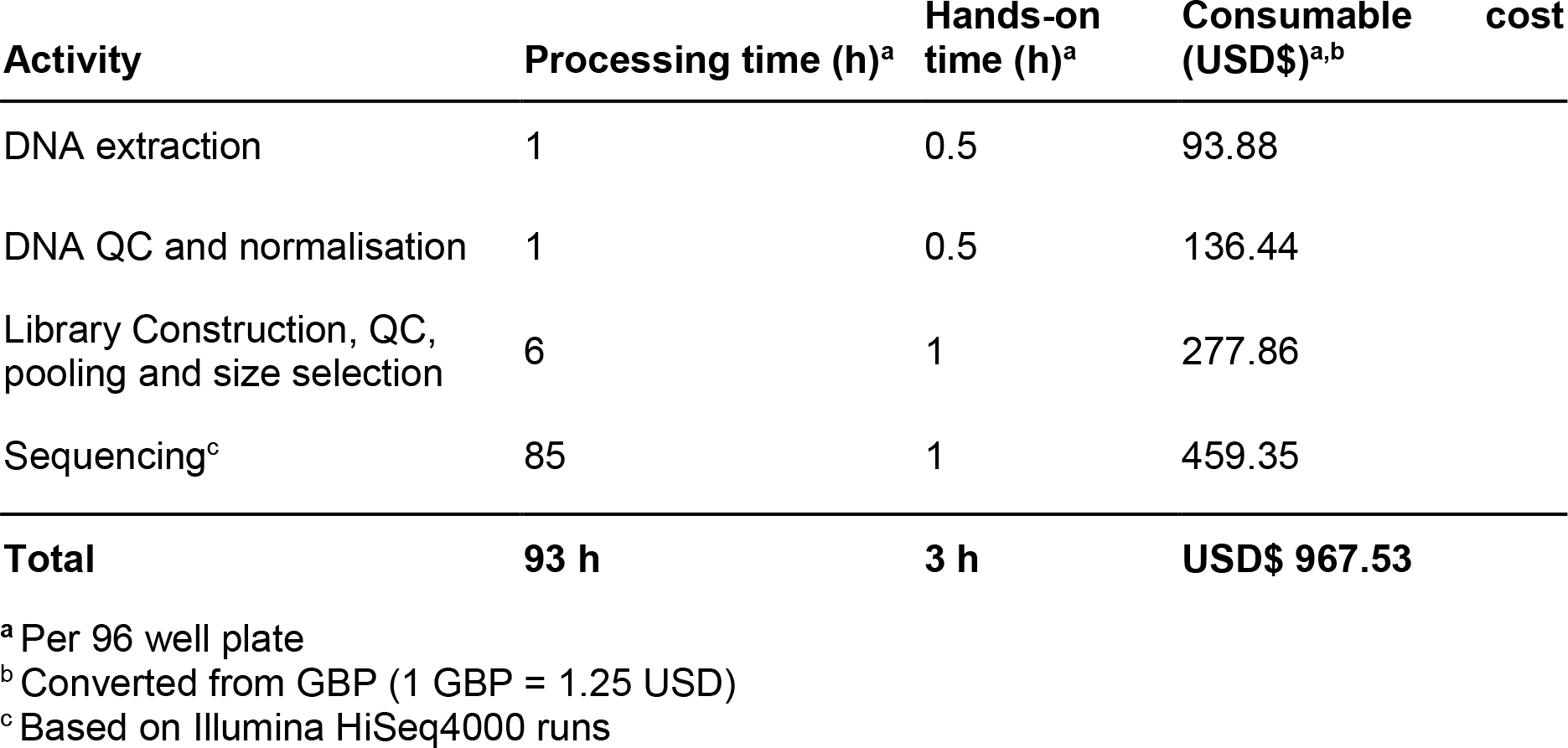
Processing time and consumable costs for DNA extraction and sequencing.

In designing the DNA extraction pipeline, we anticipated that samples would contain a wide range of DNA concentrations due to the different approaches by collaborators, some of whome sent thermolysates and others extracted DNA. The DNA was isolated in a volume of 20 μL, and the total yield ranged from 0 to 2,170 ng (average of 272 ng). Less than 6% samples contained less than 2.5 ng (Supplementary Fig. S1).

To facilitate large-scale low-cost whole-genome sequencing, we developed the LITE (Low Input, Transposase Enabled; Fig. 2) pipeline, a low-cost high-throughput library construction protocol based on the Nextera kits (Illumina). Prior to LITE library construction, all DNA samples were normalised to 0.25 ng/μL unless the concentration was below that limit, in which case samples remained undiluted. We calculated that given a bacterial genome size of 4.5 Mbp, 1 ng of DNA equated to over 200,000 bacterial genome copies. Hence the LITE pipeline was optimised to work with inputs ranging from 0.25 to 2 ng DNA. As the ratio of DNA to transposase enzyme determines the insert size of the libraries being constructed, this input amount allowed us to minimise reagent use and reaction volumes. The LITE pipeline permitted the construction of over 1,000 Illumina-compatible libraries from the 24-reaction Illumina kits, Tagment DNA Enzyme (Illumina FC 15027865) and Illumina Tagment DNA Buffer (Illumina FC 15027866).

**Fig. 2.**
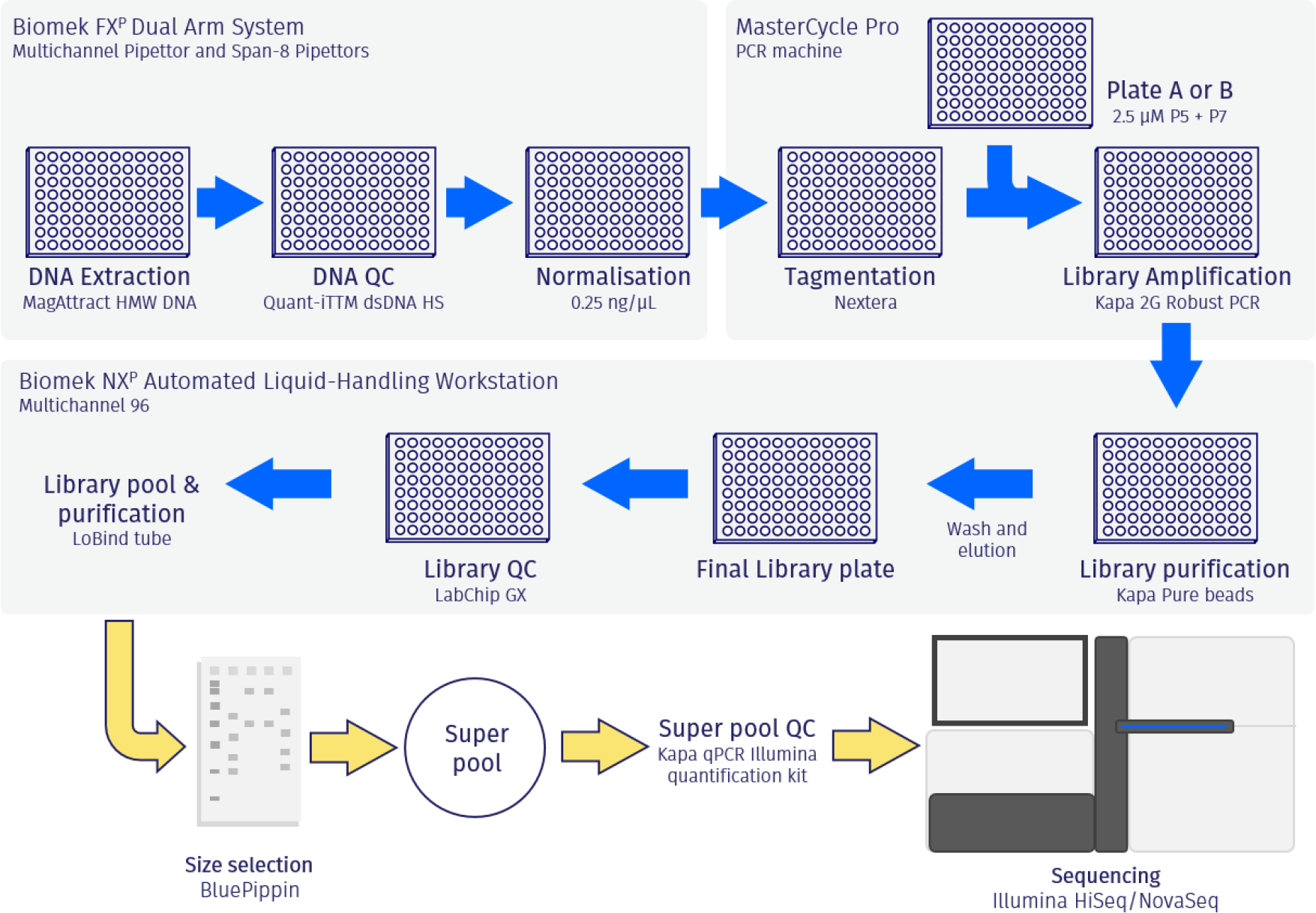
LITE (Low Input, Transposase Enabled) pipeline for library construction. The DNA was extracted using a protocol based on the MagAttract HMW DNA isolation kit (Qiagen). Library construction was performed by tagmentation using Nextera tagmentation kit, size selected on a BluePippin, and quantified using a High Sensitivity BioAnalyzer kit (Agilent) and Qubit dsDNA HS Assay (ThermoFisher). Genome sequencing of “super pools” was performed in a HiSeq™ 4000 (Illumina) system, and re-sequencing in NovaSeq™ 6000 (Illumina) when needed, both with a 2 x 150 bp paired ends read metric.

To maximise the multiplexing capability for the LITE pipeline, we designed 438 bespoke 9-bp barcodes (Supplementary Table 4), each with a hamming distance of 4 bp, giving the option to pool over 190,000 samples or uniquely dual-index more than 200 samples. The 438 barcodes allowed multiplexing capability to be maximised, and a further reduction in costs as sequencer throughputs increase in the future.

For this study we used 9-bp barcoded P7 PCR primers (Illumina) and employed twelve 6-bp barcoded P5 PCR primers (Illumina) when multiplexing 12 x 96-well plates on a HiSeq 4000 system (Illumina) and targeted a median 30x genome coverage. By using an input of only 0.5 ng DNA, combined with 14 PCR cycles consistently provided detectable amounts of library across the majority of samples.

Quality control (QC) of the resulting LITE libraries involved a Perkin Elmer LabChip® GX Nucleic Acid Analyzer. The LITE libraries typically gave three different GX electropherogram profiles depending upon whether the DNA was high molecular weight, partially degraded or completely degraded (Supplementary Fig. S2). A wide range of electropherogram profiles and the resultant molarity of library molecules was expected at this point, due to the varied approaches used by collaborators to produce and transport samples.

Up to 12 of the 96 pooled and size-selected libraries were then combined and run on a single HiSeq 4000 system lane, with a 2 x 150 bp paired-end read metric. After the initial screen was completed, samples that failed to produce 30x genome coverage were re-sequenced on a NovaSeq 6000 system, also with a 2 x 150 bp read metric. In total 1,525 (15.2%) of the 9,976 samples processed required re-sequenced, a proportion that was within the 20% contingency added to our unit cost.

### Bioinformatic analysis and data provision

To complete our WGS approach, we developed and implemented a bespoke sequence analysis bioinformatic pipeline for the *Salmonella* samples included in the study. The full pipeline is available from https://github.com/apredeus/10k_genomes. Because the estimation of sequence identity and assembly quality is relatively species-independent, and annotation is strongly species-specific, the pipeline can be easily adapted to other bacterial species by changing quality control criteria and specifying relevant databases of known proteins.

Following DNA extraction, sequencing and re-sequencing, we generated sequence reads for 9,976 (96.0%) samples, of which 7,236 were bioinformatically classified as *Salmonella enterica* using Kraken2 and Bracken^27,28^. A small proportion of the samples (209 out of 9,976; 2.1%) had been wrongly identified as *Salmonella* prior to sequencing. The remaining samples corresponded to 1,157 Gram-positive and Gram-negative bacterial isolates that were included to validate the study. The 443 (4.3%, out of the 10,419 samples received) samples that did not generate sequence reads reflected poor quality DNA extraction, due to either low biomass input or partial cell lysis. Overall, the generation of sequence data from the vast majority of samples demonstrated the robustness of the use of thermolysates coupled with the high-throughput LITE pipeline for processing thousands of samples from a variety of different collaborating organisations.

To assess the quality of sequence data, we focused on the 7,236 (69.5%) genomes identified as *Salmonella enterica* (Fig. 3). To allow the bioinformatic analysis to be customisable for other datasets, we developed a robust quality control (QC) pipeline to do simple uniform processing of all samples, and to yield the maximum amount of reliable genomic information. Well-established software tools were used to assess species-level identity from raw reads, trim the reads, assess coverage and duplication rate, assemble genomes, and to make preliminary evaluation of antibiotic resistance and virulence potential.

**Fig. 3.**
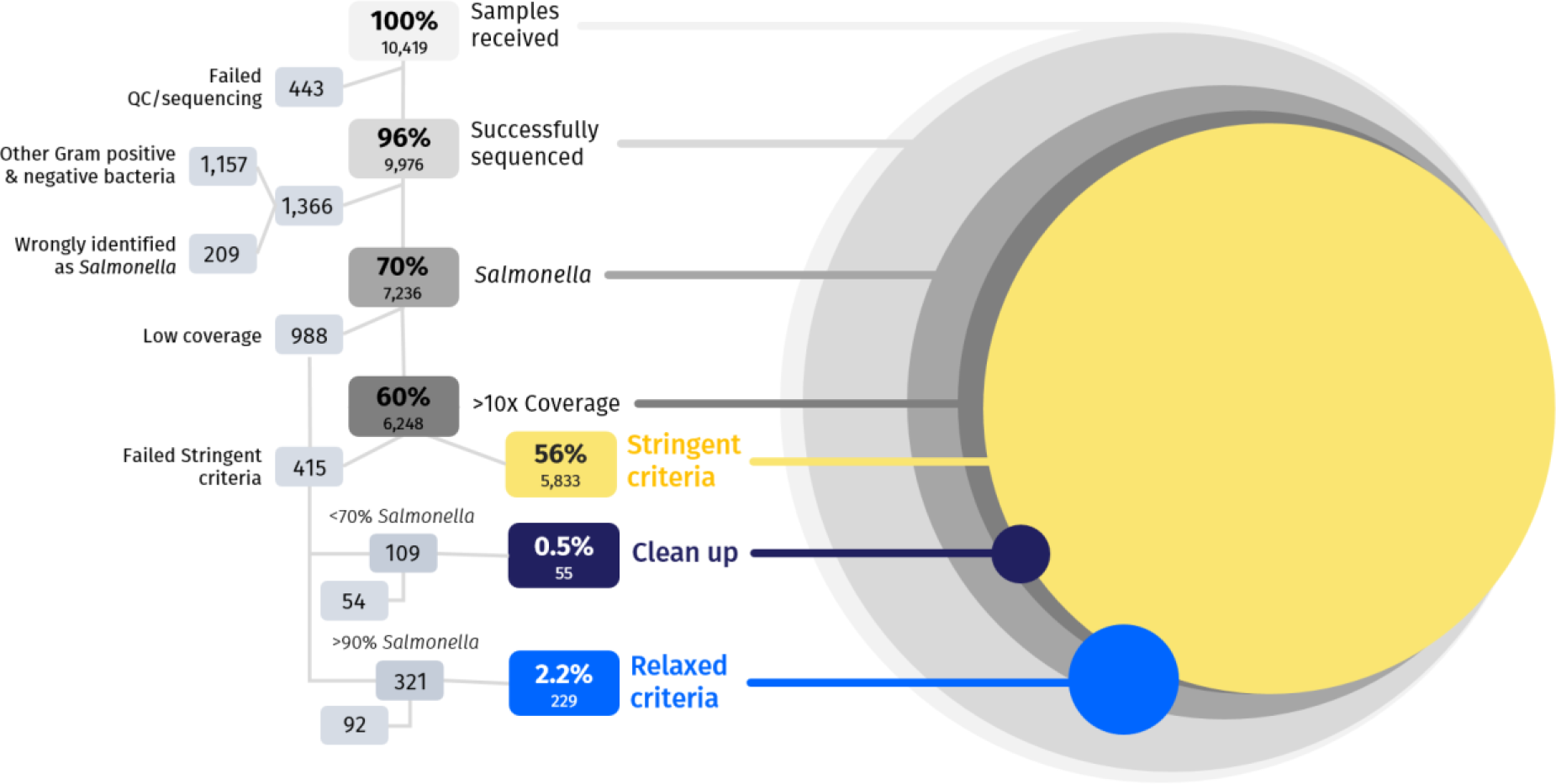
The sequential quality control process used to select whole-genome sequences for detailed analysis. Of the 10,419 isolates, 443 failed the DNA extraction or quality control prior to genome sequencing. We produced sequencing libraries of 9,975 samples, of which 1,366 were not bioinformatically-identified as *Salmonella enterica.* These 1,366 corresponded to 1,157 which were part of the 25% non-*Salmonella* component of the project, plus 209 isolates that had been mis-identified as *Salmonella* before sequencing. Of the 7,236 *Salmonella* genomes, 6,248 had sequence coverage over 10x, of which 5,833 passed the “stringent criteria”. Of the 415 samples that failed the “stringent criteria”, 284 samples were rescued based on a “clean up” (55) or a “relaxed criteria” (229). Overall, we generated 6,117 high-quality *Salmonella* genomes.

Trimming abundant adapters from the reads produced by the LITE pipeline was critical for optimal genome assembly. Using Quast^29^ and simple assembly metrics, we evaluated the performance of Trimmomatic^30^ in palindrome mode with and without retention of singleton reads, compared with BBDuk (https://jgi.doe.gov/data-and-tools/bbtools) in paired-end mode. BBDuk was selected for our analysis because this tool generated genomes with a higher N50, and a comparable number of mis-assemblies.

Genome assembly was performed using SPAdes^31^ via Unicycler^32^ in short-read mode. SPAdes is an established and widely-used tool for bacterial genome assembly, whilst Unicycler optimises SPAdes parameters and performs assembly polishing by mapping reads back to the assembled genomes. Genome assembly QC was done using the criteria established by the genome database EnteroBase^33^. Specifically, these “stringent criteria” required: 1) total assembly length between 4 and 5.8 Mb, 2) N50 of 20 kb or more, 3) fewer than 600 contigs, and 4) more than 70% sequence reads assigned to the correct species. Using this approach for *S. enterica,* 5,833 of the *Salmonella* genomes (80.6%) passed QC (Fig. 3).

To “rescue” all possible *S. enterica* in the remaining assemblies with coverage greater than 10x that failed the stringent QC, two approaches were used: “relaxed criteria” and “clean up”. The “relaxed criteria” accepted assemblies of 4 Mb to 5.8 Mb overall length, species-purity of 90% or more, N50 > 10kb, and fewer than 2,000 contigs. In contrast, the “clean up” approach was used for assemblies that had < 70% *Salmonella* sequence reads using the “stringent criteria”. The raw reads of these samples were “cleaned” using Kraken2 & Bracken, with the reads assigned to *Salmonella* being retained, and subjected to the “stringent criteria” for QC detailed above. The assemblies rescued by these two approaches accounted for a further 3.9% (284) assemblies from our initial *Salmonella* collection. In total, we generated 6,117 high quality *S. enterica* genomes, corresponding to 84.5% of the total *Salmonella* isolates successfully sequenced through the LITE pipeline (Fig. 3 and 4).

**Fig. 4.**
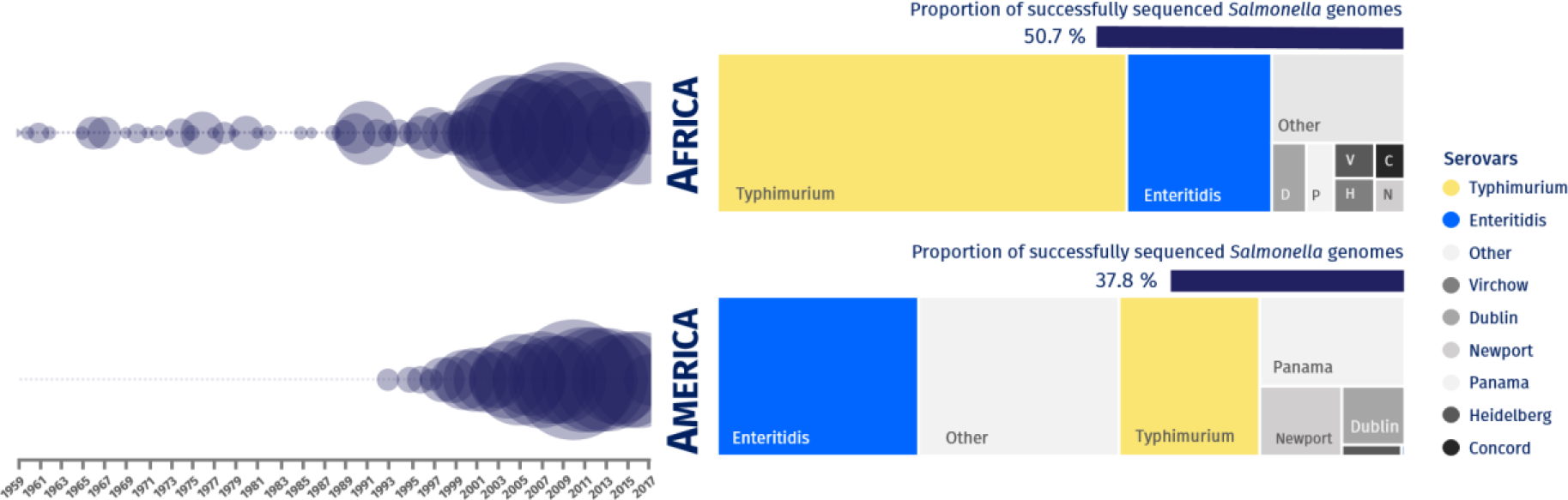
Genome-based summary of *Salmonella enterica* from African and American datasets, organised by continent, year of isolation, and serovar. Of the 6,117 *Salmonella enterica* genomes that were successfully sequenced and that passed QC, 3,100 (50.7%) were from Africa and 2,313 (37.8%) were from America. Bubble size represents the number of genomes isolated between 1959 and 2017. The graphs represent the proportion of the main *Salmonella* serovars predicted based on genome analysis: 1,844 *S.* Typhimurium & 657 *S.* Enteritidis from Africa, and 474 *S.* Typhimurium & 676 *S.* Enteritidis from America.

Genome sequence data were shared with collaborators via downloadable packages hosted by the Centre of Genomic Research, University of Liverpool (UK). These packages included sequencing statistics, raw (untrimmed) fastq files of sequence reads, and the individual genome assemblies. We included the genome-derived *Salmonella* serovar and sequence type of each isolate (Fig. 4).

Together with predicted sequence type and serovar, the genome-derived information was provided to permit local surveillance laboratories and infectious disease clinicians to derive important insights about the *Salmonella* variants circulating in their countries. The value of bacterial WGS data for generating epidemiological insights or understanding pathogen evolution has been summarised recently^19^. All the processed sequence reads and assemblies were deposited in the European Nucleotide Archive under the project accession number PRJEB35182 (ERP118197). Individual accession numbers are listed in Supplementary Table 3.

## Discussion

We have optimised an efficient and relatively inexpensive method for large-scale collection and sequencing of bacterial genomes, by streamlining the collection of isolates, and developing a logistics pipeline that permitted ambient shipment of thermolysates. The global focus of our study provided a diverse collection of 10,419 clinical and environmental bacterial isolates for a single sequencing study within one year.

The effectiveness and accessibility of our approach allowed all samples to be collected in a timely manner, and generated genomic data for LMI countries that lacked easy access to sequencing technology. The novel optimised DNA extraction and sequencing LITE pipeline allowed bacterial genomes to be generated at a consumables cost of USD$10 per sample (the full economic cost cannot be calculated because collaborator staff time was an in-kind contribution). This optimised DNA extraction and sequencing pipeline, in conjunction with the generation of thermolysates, provides a robust approach for global collaboration on the genome-based mass surveillance of pathogens.

However, our approach did pose manual and logistical challenges. We propose that for future implementations of a similar approach for sequencing thousands of bacterial isolates, it is important to make an early investment in the development of a shared, protected and version controlled database to store epidemiological information, coupled with automated scripts to handle sequencing data, and a streamlined system for the sending and receiving of samples.

Our method is suitable for other large collections of Gram-negative or Gram-positive bacteria, and is designed to complete an academic genome sequencing project within a limited time-frame (one year). However, the LITE pipeline represents a compromise in terms of data quality to maximise economic value. It is important that all QC steps and the rigorous bioinformatic approach that we specify are followed to produce a reliable dataset, which in this case generated 84.5% high-quality genomes of the 7,236 successfully-sequenced *Salmonella* isolates (Fig. 3 and 4).

A key aspect of our methodology was the involvement of researchers fluent in multiple languages, to maximise clear communication and ensure access to countries across the world. The approach will be particularly relevant when rapid, low-cost, and collaborative genome sequencing of bacterial pathogens is required. Our concerted approach demonstrates the value of true global collaboration, offering potential for tackling international epidemics or pandemics in the future.

## Acknowledgements

The project was supported by both a Global Challenges Research Fund (GCRF) data & resources grant BBS/OS/GC/000009D and the BBSRC Core Capability Grant to the Earlham Institute BB/CCG1720/1. Next-generation sequencing and library construction were delivered via the BBSRC National Capability in Genomics and Single Cell (BB/CCG1720/1) at Earlham Institute, by members of the Genomics Pipelines Group. Each of the following Earlham Institute staff and alumni made an enormous contribution to the project: Dr Helen Chapman, Mr. Jake Collins, and Miss Sophie Stephenson. This project was partly supported by the Wellcome Trust Senior Investigator Award (106914/Z/15/Z) to Jay C. D. Hinton. We are grateful to the Centre for Genomic Research, University of Liverpool (UK) for computing support.

## Author contributions

NH and JCDH conceived the idea and received funding. BPS wrote the manuscript. JCDH, NH, CVP, AVP, DH, BK, KSB, WR and NAF contributed to manuscript writing and editing. The 10KSG consortium reviewed the manuscript. JCDH, NH, NAF, KSB, CVP and BPS designed the study. BPS and KC curated the metadata. BPS was the main point of contact for the 10KSG consortium, designed and prepared protocols & other material for collaborators, and distributed barcoded tubes. WR and BPS designed web page. RL uploaded generated data to ENA. CW and NS supervised logistics at the Earlham Institute. BPS, CVP and HW optimised thermolysates generation. AVP, CVP, RL and CS developed bioinformatic pipelines and analysis. DH and JL optimised LITE protocol. BPS, CVP and the 10KSG consortium isolated and prepared bacterial samples.

## The 10,000 *Salmonella* genomes (10KSG) Consortium (in alphabetical order)

Blanca M. Perez-Sepulveda^1^, Darren Heavens^2^, Caisey V. Pulford^1^ & María Teresa Acuña^11^, Dragan Antic^1^, Martin Antonio^5^, Kate S. Baker^1^, Johan Bernal^8^, Hilda Bolaños^11^, Marie Chattaway^9^, Angeziwa Chirambo^4^, Karl Costigan^1^, Saffiatou Darboe^5^, Paula Díaz^10^, Pilar Donado^8^, Carolina Duarte^10^, Francisco Duarte^11^, Dean Everett^4^, Séamus Fanning^12^, Nicholas A. Feasey^3,4^, Patrick Feglo^13^, Adriano M. Ferreira^15^, Rachel Floyd^1^, Ronnie G Gavilán^13,26^, Melita A. Gordon^1,4^, Neil Hall^2^, Rodrigo T. Hernandes^15^, Gabriela Hernández-Mora^16^, Jay C. D. Hinton^1^, Daniel Hurley^12^, Irene N. Kasumba^17^, Benjamin Kumwenda^7^, Brenda Kwambana-Adams^24^, James Lipscombe^2^, Ross Low^2^, Salim Mattar^18^, Lucy Angeline Montaño^10^, Cristiano Gallina Moreira^15^, Jaime Moreno^10^, Dechamma Mundanda Muthappa^12^, Satheesh Nair^9^, Chris M. Parry^3^, Chikondi Peno^4^, Jasnehta Permala-Booth^17^, Jelena Petrović^19^, Alexander V. Predeus^1^, José Luis Puente^20^, Getenet Rebrie^21^, Martha Redway^1^, Will Rowe^1,6^, Terue Sadatsune^15^, Christian Schudoma^2^, Neil Shearer^2^, Claudia Silva^20^, Anthony M. Smith^22,25^, Sharon Tennant^17^, Alicia Tran-Dien^23^, Chris Watkins^2^, Hermione Webster^1^, François-Xavier Weill^23^, Magdalena Wiesner^10^, Catherine Wilson^1,4^

^1^IVES, University of Liverpool, Liverpool, UK

^2^Earlham Institute, Norwich Research Park, Norwich, UK

^3^Liverpool School of Tropical Medicine, Liverpool, UK

^4^Malawi-Liverpool-Wellcome Programme, Blantyre, Malawi

^5^Medical Research Council Unit The Gambia at LSHTM

^6^University of Birmingham, Birmingham, UK

^7^College of Medicine, University of Malawi, Blantyre, Malawi

^8^Corporación Colombiana de investigación Agropecuaria AGROSAVIA, Colombia

^9^Public Health England, UK

^10^Instituto Nacional de Salud (INS), Colombia

^11^Instituto Costarricense de Investigación y Enseñanza en Nutrición y Salud (INCIENSA), Costa Rica

^12^University College Dublin, Ireland

^13^Kwame Nkrumah University of Science and Technology, Ghana

^14^Instituto Nacional de Salud, Lima, Peru

^15^São Paulo State University (UNESP), Brazil

^16^Bacteriology Laboratory, Servicio Nacional de Salud Animal (SENASA), Costa Rica

^17^University of Maryland School of Medicine, USA

^18^Instituto de Investigaciones Biológicas del Trópico, Universidad de Córdoba, Colombia

^19^Scientific Veterinary Institute Novi Sad, Serbia

^20^Instituto de Biotecnología, UNAM, Mexico

^21^University of Jimma, Ethiopia

^22^National Institute for Communicable Diseases (NICD), South Africa

^23^Institut Pasteur, Paris, France

^24^University College London, London, UK

^25^University of the Witwatersrand, South Africa

^26^Escuela Profesional de Medicina Humana, Universidad Privada San Juan Bautista, Lima, Peru

## Supplementary material

**Supplementary Table 1.** Metadata template form

**Supplementary Table 2.** Optimisation of bacterial thermolysates generation and DNA extraction.

**Supplementary Table 3.** Metadata for sequenced isolates, including bioinformatic stats for *Salmonella* genomes and ENA accession numbers.

**Supplementary Table 4.** Bespoke 9 bp barcodes for library construction using the LITE pipeline

**Supplementary figure S1:**
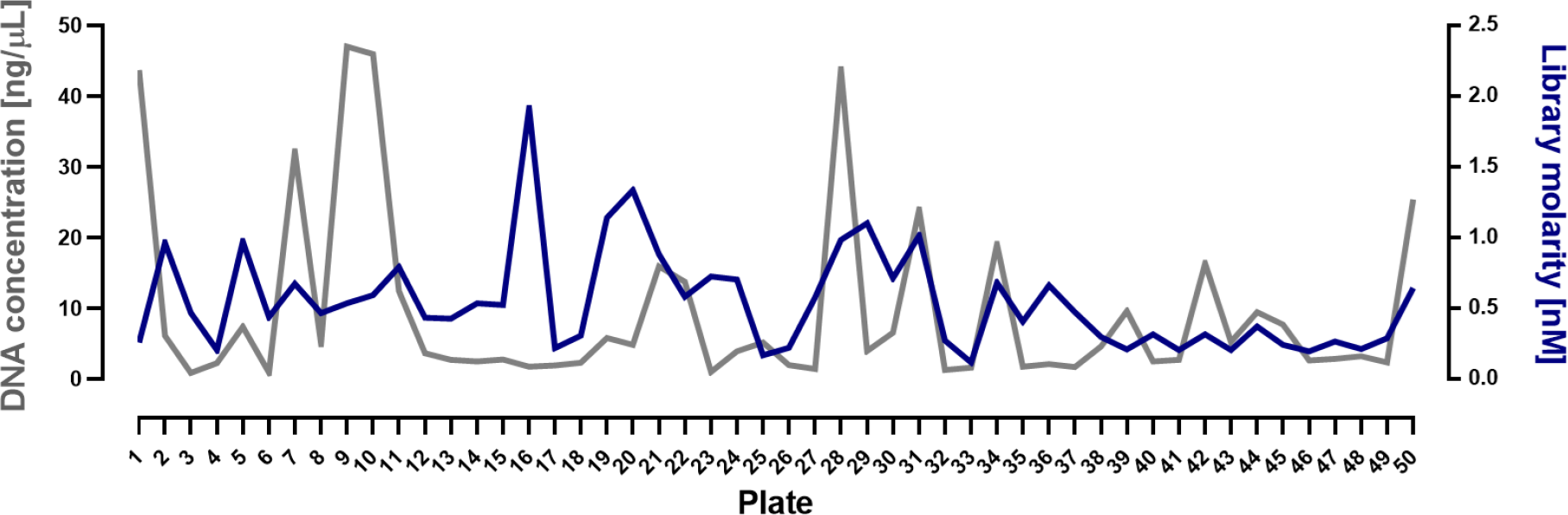
Average DNA concentration and molarity of libraries constructed using the LITE pipeline across individual 96-well plates. Average DNA concentrations (grey) and library molarity between 400 and 600 bp (blue) are shown for the first fifty 96-well plates that were processed.

**Supplementary figure S2:**
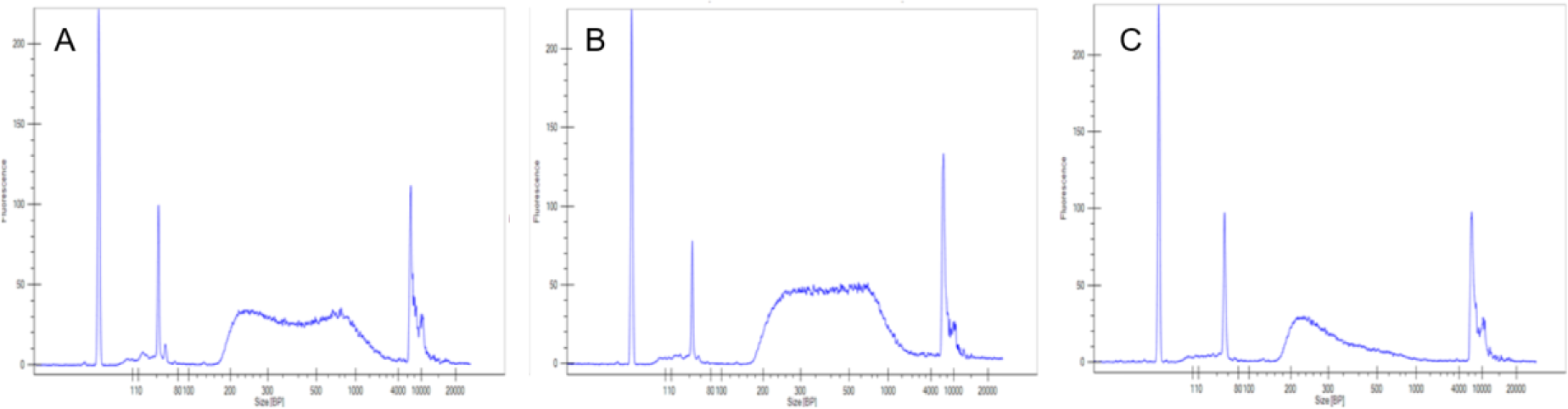
Assessment of DNA integrity amongst libraries constructed using the LITE pipeline. Perkin Elmer GX electropherograms of exemplar LITE libraries: (A) high quality HMW DNA, (B) partially-degraded DNA, and (C) degraded DNA.

## Methods

### Study design and optimisation

We designed the project with the aim of validating an efficient method for large-scale assembly and sequencing of bacterial genomes. We selected *Salmonella* as a model organism due to its worldwide relevance and current burden of infection. We aimed to assemble a pool of bacterial samples that would represent the different scenarios, including a 25% of non-*Salmonella* isolates, to allow the method to be extrapolated to other bacterial datasets. The 25% of non-*Salmonella* organisms were selected to cover Gram-negative *(Shigella* and *Klebsiella)* and Gram-positive *(Staphylococcus)* bacteria. The targeted *Salmonella* isolates were predominantly *S.* Enteritidis and Typhimurium, and associated with human bloodstream infection. However we expanded the sampling criteria to other serovars, body compartments and source types to include some animal and environmental samples.

Method optimisation focused on standardising a safe protocol for sample transport and processing. Briefly, the optimised method comprised bacterial isolates grown at 37°C overnight directly in FluidX tubes (FluidX tri-coded jacket 0.7 mL, 68-0702-11, Brooks Life Sciences) with 100 μL rich media (LB or Buffered Peptone) from a frozen stock (one “scoop” or bead (Microbank™, Pro Lab Diagnostics Inc.). Then, the samples were inactivated by incubation at > 95°C for 20 min, followed by storage at 4°C until collection. Sample transportation was carried out at ambient temperature.

We optimised this method using *Salmonella enterica* serovar Typhimurium D23580, *Eschericchia coli* K12, and *Staphylococcus aureus* Newman, selecting either a “scoop” with a 10 μL plastic loop taken from a bacterial glycerol (50% v/v) stock or 2 beads of bacteria stored at −80°C in a Microbank tube™ cryotubes (Pro-Lab Diagnostics). The samples were grown at 37°C and 220 rpm overnight in either 100 or 200 μL LB (1% tryptone, 0.5% yeast extract, 0.5% NaCl; pH 7.0). 100 μL of each sample was heated to either 90°C, 95°C or 100°C for 10 min or 20 min, and then plated on nutrient agar (1.5% Agar-LB) for CFU determination (Supplementary Table 2).

To test the effect of transport, the samples were subjected to genomic DNA extraction using a DNeasy Blood & Tissue Kit (Qiagen) after incubation at room temperature for more than 7 days. The quality of extracted DNA was assessed by 1% agarose gel electrophoresis, and fluorometric DNA quantification using Qubit™ dsDNA HS Assay Kit (Invitrogen™) (Supplementary Table 2).

Detailed protocols were sent to collaborators, along with a metadata template and barcoded tubes. The design of the metadata template and protocol booklet was tested several times for clarity and to obtain unified information avoiding different interpretations by the user. The metadata template (Supplementary Table 1) was a Microsoft Excel spreadsheet divided in five main categories: 1) Unique identifiers, with information about pre-read barcodes, including plate & tube barcode, tube location, and replacement barcode, 2) Isolate details, encompassing information about strain name, bacterial species & serovar (*Salmonella* only), sender, date and location of isolation, and type of sample submitted (DNA, thermolysates or preserved culture), 3) Sample type, with detailed information about source of isolation, such as human, animal or environmental origin, and 4) Antimicrobial resistance phenotype of tested antimicrobials. We also added an extra column for relevant information that could not be assigned to any other category, such as type of study and references. The metadata collected were stored per collaborator and then combined into a metadata master form for curation. Curation was done manually, standardising each category by column and keeping version control. The final metadata master form was cross-referenced with the list of sent barcodes for inconsistencies.

### DNA extraction and normalisation

DNA was extracted from bacterial thermolysates on a Biomek FX^P^ instrument using a protocol based on the MagAttract HMW DNA isolation kit (Qiagen). Incomplete barcoded 96-tube plates received were re-organised and FluidX barcodes re-read using the FluidX barcode reader and software prior to DNA extraction, to determine plate layouts. The tubes were de-capped using a manual eight-tube decapper and the cellular material was re-suspended using a multichannel pipette. Up to 100 μL of the suspension were transferred to a clean 96-well plate. The plate was spun at 4,000 rpm in an Eppendorf 5810R centrifuge to pellet the cells and discard the supernatant.

Cell pellets were re-suspended in a mixture of 12 μL of Qiagen ATL buffer and 2 μL Proteinase K, and incubated at 56°C for 30 min in an Eppendorf Thermomixer C. The samples were cooled to room temperature, and 1 μL of MagAttract Suspension G was added. The samples were mixed, and 18.67 μL of Qiagen MB buffer were added, followed by mixing. The samples were incubated for 3 min and placed on a 96-well magnetic particle concentrator (MPC) to pellet the beads. The supernatant was discarded, and whilst remaining on the MPC the beads were washed once with 45 μL Qiagen MW1 buffer and once with 45 μL Qiagen PE buffer. The recommended water washes were omitted to help increase yield.

The plate was then removed from the MPC and, using a new set of filter tips, 20 μL of Qiagen AE buffer wae added and the samples mixed to re-suspend the beads. The samples were incubated at room temperature for 3 min to elute the DNA. The plate was placed back on the MPC and the DNA was transferred to a new 96-well plate.

The concentration of each sample was determined using the Quant-iT™ dsDNA Assay, high sensitivity kit (ThermoFisher). A standard curve was generated by mixing 10 μL of the eight DNA standards provided (0 to 10 ng/μL) with 189 μL of 1x Quant-iTTM dsDNA HS buffer, 1 μL of Quant-iTTM dsDNA HS reagent and 1 μL of DNA in a 96-well black Greiner plate. The fluorescence was detected on a Tecan Infinite F200 Pro plate reader (Tecan).

For samples received as DNA, 198 μL of 1x Quant-iTTM dsDNA HS buffer, 1 μL of Quant-iTTM dsDNA HS reagent and 1 μL of DNA were combined in a 96-well black Greiner plate, and the fluorescence detected using the Tecan plate reader. Concentrations were calculated using the standard curve, and the DNA was normalised to 0.25 ng/μL in elution buffer using the Biomek FX^P^ instrument.

### Library construction and sequencing

A master mix containing 0.9 μL of Nextera buffer, 0.1 μL Nextera enzyme and 2 μL of DNAse free water was combined with 2 μL of normalised DNA. This reaction was incubated at 56°C for 10 min on an Eppendorf MasterCycle Pro PCR instrument. 2 μL of an appropriately barcoded 2.5 μM P7 adapter were added, and then 18 μL of a master mix containing 2 μL of an appropriately barcoded 2.5 μM P5, 5 μL Kapa Robust 2G 5x reaction buffer, 0.5 μL 10 mM dNTPs, 0.1 μL Kapa Robust 2G polymerase and 10.4 μL DNase free water were added to the tube. This reaction was then subjected to PCR amplification as follows: 72°C x 3 min, 98°C for 2 min, then 14 cycles of 98°C x 10 s, 62°C x 30 s and 72°C x 3 min, followed by a final incubation at 72°C for 5 min on an Eppendorf MasterCycle Pro.

The amplified library was then subjected to a magnetic bead-based purification step on a Biomek NX^P^ instrument. 25 μL of Kapa Pure beads (Roche, UK) were added to 25 μL of amplified library, and mixed. This library was incubated at room temperature for 5 min, briefly spun in an Eppendorf 5810R centrifuge and placed on a 96-well magnetic particle concentrator. Once the beads had pelleted, the supernatant was removed and discarded, and the beads washed twice with 40 μL of freshly prepared 70% ethanol. After the second ethanol wash, the beads were left to air dry for 5 min. The 96-well plate was removed from the MPC and the beads were re-suspended in 25 μL of 10 mM TRIS-HCl, pH 8 (Elution Buffer). The DNA was eluted by incubating the beads for 5 min at room temperature. The plate was replaced on the MPC, the beads allowed to pellet, and the supernatant containing the DNA was transferred to a new 96-well plate.

To assess the concentrations of individual libraries, 20 μL of elution buffer was added to 2 μL of purified library, and run on a LabChip GX (Perking Elmer) using the High throughput, High Sense reagent kit and HT DNA Extended Range Chip according to manufacturers’ instructions. To determine the amount of material present in each library between 400 and 600 bp, a smear analysis was performed using the GX analysis software. The resulting value was used to calculate the amount of each library to pool. Pooling of each 96-libraries was performed using a Biomek Nx instrument. 100 μL of the pooled libraries were added to 100 μL of Kapa Pure beads in a 1.5 mL LoBind tube. The sample was vortexed and incubated at room temperature for 5 min to precipitate the DNA onto the beads. The tube was then placed on an MPC to pellet the beads, the supernatant discarded, and the beads were washed twice with 200 μL of freshly prepared 70% ethanol. The beads were left to air dry for 5 min and then re-suspended in 30 μL Elution Buffer. The samples were incubated at room temperature for 5 min to elute the DNA. The plate was placed back on the MPC and the DNA was transferred to a new 1.5 mL tube.

The concentrated sample containing a pool of 96 libraries was subjected to size selection on a BluePippin (Sage Science, Beverly, USA). The 40 μL in each collection well of a 1.5% BluePippin cassette were replaced with fresh running buffer, and the separation and elution current checked prior to loading the sample. 10 μL of R2 marker solution were added to 30 μL of the pooled library, and then the combined mixture was loaded into the appropriate well.

Using the smear analysis feature of Perkin Elmer GX software, we calculated the amount of material between 400 and 600 bp for each library. We targeted this region based on the electropherograms in Supplementary Fig. S2, to minimise the overlap between 150 bp paired end reads and maximise the number of libraries that would generate data. We determined the detection limit for the molarity within this size range to be 0.007 nM, meaning that libraries with lower concentrations were reported as 0.007 nM. The amount of library material between 400 to 600 bp ranged from 0.0 to 2.4 nM (average of 0.3 nM), with less than 6% having less than 0.007 nM (Supplementary Fig. S1).

Post size selection, the 40 μL from the collection well were recovered, and the library size was determined using a High Sensitivity BioAnalyzer kit (Agilent) and DNA concentration calculated using a Qubit dsDNA HS Assay (ThermoFisher). “Super pools” were created by equimolar pooling of up to 12 size-selected 96-sample pools, each with a different P5 barcode. Using these molarity figures, 96 libraries were equimolarly-pooled, concentrated and then size-selected using a 1.5% cassette on the Sage Science Blue Pippin.

To determine the number of viable library molecules, the super pools were quantified using the Kapa qPCR Illumina quantification kit (Kapa Biosystems) prior to sequencing. For the initial screen, sequencing was performed on the HiSeq™ 4000 (Illumina). For re-sequencing of samples, the sequencing was carried out in a lane of an S1 flowcell on the NovaSeq™ 6000 (Illumina), both with a 2×150 bp read metric.

### Bioinformatic analysis and data distribution

Raw sequencing reads (paired-end, 2×150 bp) were examined using FastQC v0.11.8 (https://www.bioinformatics.babraham.ac.uk/projects/fastqc), confirming 0-20% Nextera adapter sequence presence in all examined reads. Quick coverage estimation was done raw unaligned reads, assuming genome length of 4.8 Mb for *Salmonella enterica*. Taxonomic classification of raw reads was performed using Kraken v2.0.8-beta^1^ with Minikraken 8GB 201904_UPDATE database, followed by species-level abundance estimation using Bracken v1.0.0^2^ with distribution for 150 bp k-mer. Sequence duplication level was estimated by alignment of reads using Bowtie v2.3.5^3^ to genome assembly of LT2 strain (NCBI accession number GCA_000006945.2), followed by MarkDuplicates utility from Picard tools v2.21.1 (http://broadinstitute.github.io/picard).

Raw sequence reads were then trimmed and assembled using Uncycler v0.4.7^4^ in short-read mode. Several trimming strategies were tested including quality trimming with seqtk (https://github.com/lh3/seqtk) followed by Trimmomatic v0.39^5^ in palindromic mode with and without retaining the single reads, and BBDuk v38.07 (https://jgi.doe.gov/data-and-tools/bbtools). We evaluated the resulting assemblies using overall length, N50, and number of contigs. Genome assembly quality was done using a the criteria established on EnteroBase^6^ (https://enterobase.readthedocs.io/en/latest) for *S. enterica:* 1) total assembly length between 4 and 5.8 Mb; 2) N50 of 20 kb or more; 3) fewer than 600 contigs; 4) more than 70% correct species assigned by Kraken (in our case, the latter was replaced with Kraken2+Bracken assessment of the raw reads). Samples that failed the stringent criteria were divided into two groups. Group 1 were subjected to “relaxed criteria”, which included assemblies of 4 Mb - 5.8 Mb overall length, species purity of 90% or more, N50 >10,000, and fewer than 2,000 contigs. Group 2 included samples that had less than 70% *Salmonella* by original assessment, but produced assemblies passing the stringent criteria from “cleaned up” reads obtained by keeping only raw reads assigned *S. enterica* by Kraken2 + Bracken.

Assembled *Salmonella* genomes were annotated using Prokka v1.13.7^7^ using a a custom protein database generated from *S. enterica* pan-genome analysis. Additionally, *Salmonella* assemblies were *in silico* serotyped using command line SISTR v1.0.2^8^ and assigned sequence type using mlst v2.11^9^ (https://github.com/tseemann/mlst). We have used cgMLST serovar assignment provided by SISTR for all further classification and comparison with metadata. Preliminary resistance and virulence gene profiling was done using Abricate v0.9.8 (https://github.com/tseemann/abricate). All processing scripts detailing command settings and custom datasets are available at https://github.com/apredeus/10k_genomes.

Data distribution was carried out by sharing packages through links created at the Centre for Genomic Research, University of Liverpool (UK). The packages contained sequencing stats, raw (untrimmed) fastq read files, assemblies, and a text files with information about serovar and sequence type details. All the processed reads and assemblies were deposited in the European Nucleotide Archive using the online portal Collaborative Open Plant Omics (COPO; https://copo-project.org/copo) under the project accession number PRJEB35182 (ERP118197). COPO is an online portal for the description, storage and submission of publication data. The COPO wizards allow users to describe their data using ontologies to link and suggest metadata to include based on past submissions and similar projects. This enables meaningful description and therefore easy retrieval of the data in addition to standardising the format, thereby removing most of the hassle from data submission. Individual accession numbers are listed in Supplementary Table 3.

### Code availability

Our code is available as open source (GPL v3 license) at https://github.com/apredeus/10k_genomes

### Data availability

All sequencing datasets used in this study are publicly available in the European Nucleotide Archive under the project accession number PRJEB35182 (ERP118197). Individual accession numbers are listed in Supplementary Table 3.

